# Topographic Divergence of Atypical Cortical Asymmetry and Regional Atrophy Patterns in Temporal Lobe Epilepsy: A Worldwide ENIGMA Study

**DOI:** 10.1101/2021.04.30.442117

**Authors:** Bo-yong Park, Sara Larivière, Raul Rodríguez-Cruces, Jessica Royer, Shahin Tavakol, Yezhou Wang, Lorenzo Caciagli, Maria Eugenia Caligiuri, Antonio Gambardella, Luis Concha, Simon S. Keller, Fernando Cendes, Marina K. M. Alvim, Clarissa Yasuda, Leonardo Bonilha, Ezequiel Gleichgerrcht, Niels K. Focke, Barbara A. K. Kreilkamp, Martin Domin, Felix von Podewils, Soenke Langner, Christian Rummel, Michael Rebsamen, Roland Wiest, Pascal Martin, Raviteja Kotikalapudi, Benjamin Bender, Terence J. O’Brien, Meng Law, Benjamin Sinclair, Lucy Vivash, Patricia M. Desmond, Charles B. Malpas, Elaine Lui, Saud Alhusaini, Colin P. Doherty, Gianpiero L. Cavalleri, Norman Delanty, Reetta Kälviäinen, Graeme D. Jackson, Magdalena Kowalczyk, Mario Mascalchi, Mira Semmelroch, Rhys H. Thomas, Hamid Soltanian-Zadeh, Esmaeil Davoodi-Bojd, Junsong Zhang, Matteo Lenge, Renzo Guerrini, Emanuele Bartolini, Khalid Hamandi, Sonya Foley, Bernd Weber, Chantal Depondt, Julie Absil, Sarah J. A. Carr, Eugenio Abela, Mark P. Richardson, Orrin Devinsky, Mariasavina Severino, Pasquale Striano, Costanza Parodi, Domenico Tortora, Sean N. Hatton, Sjoerd B. Vos, John S. Duncan, Marian Galovic, Christopher D. Whelan, Núria Bargalló, Jose Pariente, Estefania Conde, Anna Elisabetta Vaudano, Manuela Tondelli, Stefano Meletti, Xiang-Zhen Kong, Clyde Francks, Simon E. Fisher, Angelo Labate, Sanjay M. Sisodiya, Paul M. Thompson, Carrie R. McDonald, Andrea Bernasconi, Neda Bernasconi, Boris C. Bernhardt

## Abstract

Temporal lobe epilepsy (TLE), a common drug-resistant epilepsy in adults, is primarily a limbic network disorder associated with predominant unilateral hippocampal pathology. Structural MRI has provided an *in vivo* window into whole-brain grey matter pathology in TLE relative to controls, by either mapping (i) atypical inter-hemispheric asymmetry or (ii) regional atrophy. However, similarities and differences of both atypical asymmetry and regional atrophy measures have not been systematically investigated. Here, we addressed this gap using the multi-site ENIGMA-Epilepsy dataset comprising MRI brain morphological measures in 732 TLE patients and 1,418 healthy controls. We compared spatial distributions of grey matter asymmetry and atrophy in TLE, contextualized their topographies relative to spatial gradients in cortical microstructure and functional connectivity, and examined clinical associations using machine learning. We identified a marked divergence in the spatial distribution of atypical inter-hemispheric asymmetry and regional atrophy mapping. The former revealed a temporo-limbic disease signature while the latter showed diffuse and bilateral patterns. Our findings were robust across individual sites and patients. Cortical atrophy was significantly correlated with disease duration and age at seizure onset, while degrees of asymmetry did not show a significant relationship to these clinical variables. Our findings highlight that the mapping of atypical inter-hemispheric asymmetry and regional atrophy tap into two complementary aspects of TLE-related pathology, with the former revealing primary substrates in ipsilateral limbic circuits and the latter capturing bilateral disease effects. These findings refine our notion of the neuropathology of TLE and may inform future discovery and validation of complementary MRI biomarkers in TLE.

## Introduction

Temporal lobe epilepsy (TLE) is the most common drug-resistant epilepsy in adults. Its hallmark is pathology of mesiotemporal structures, notably the hippocampus, entorhinal cortex, amygdala, and temporal pole (Falconer *et al*., 1964; Margerison and Corsellis, 1966; Blanc *et al*., 2011; Blümcke *et al*., 2013; Thom, 2014). The degree of atrophy in these regions correlates with the tendency to express epileptic activity (Bartolomei *et al*., 2005; Ogren *et al*., 2009). Moreover, unilateral anteromesial resection leads to worthwhile improvement in approximately 90% of patients and long-term seizure freedom in more than 50% (Wiebe *et al*., 2001; de Tisi *et al*., 2011; Bernhardt *et al*., 2015).

Magnetic resonance imaging (MRI) can identify the pathological substrate of TLE *in vivo*, and lateralize and define the surgical target. Indeed, MRI has provided biomarker candidates for TLE diagnostics, prognostics, and disease staging (Bernhardt *et al*., 2013*a*; Winston *et al*., 2013, Larivière *et al*., 2020*a*). MRI analyses in TLE traditionally focus on manually or automatically delineating individual mesiotemporal structures, followed by (i) the analysis of inter-hemispheric grey matter asymmetry or (ii) the regional comparison of morphometric measures in patients relative to healthy controls. Studies focusing on mesiotemporal grey matter consistently reported atrophy and marked asymmetry, reaffirming that TLE is primarily a limbic disorder (Cascino *et al*., 1991; Cendes *et al*., 1993; van Paesschen *et al*., 1995; Kuzniecky *et al*., 1997; Briellmann *et al*., 1998; Bernasconi *et al*., 2003; Bonilha *et al*., 2004, 2007; Bernhardt *et al*., 2015).

With advancements and automation of image processing techniques, quantitative MRI analysis has been extended to the whole-brain level using volumetric analysis and voxel-based morphometry (Bonilha *et al*., 2004, 2005, 2009, 2010; Seidenberg *et al*., 2005; Pulsipher *et al*., 2007; Keller and Roberts, 2008) as well as surface-based cortical thickness analysis (Lin *et al*., 2007; Bernhardt *et al*., 2008, 2010; McDonald *et al*., 2008; Galovic *et al*., 2019). These studies have mainly been cross-sectional regional comparisons between TLE and healthy controls, and explored patterns of inter-hemispheric asymmetry in TLE only sporadically. Whole-brain analyses often showed asymmetric mesiotemporal damage, and also revealed widespread and bilateral decreases in cortical grey matter outside the mesiotemporal lobe, with neither a limbic nor lateralized predominance (McDonald *et al*., 2008, Bernhardt *et al*., 2009*a*; Whelan *et al*., 2018; Galovic *et al*., 2019; Weng *et al*., 2020). Similar findings were confirmed by a multi-site initiative aggregating and analyzing brain morphometric measures in common epilepsies (Whelan *et al*., 2018; Sisodiya *et al*., 2020).

Outside the mesiotemporal regions, the scarcity of asymmetry analyses precludes insights into how similar or different patterns of atypical structural asymmetry are relative to patterns of regional atrophy in TLE. Analyzing both atrophy and asymmetry features could inform the development of individualized MRI biomarkers (Bernasconi *et al*., 2003; Bonilha *et al*., 2003, 2004; Alhusaini *et al*., 2012). Moreover, comparing these patterns could clarify whether these reflect different disease processes. One emerging family of approaches stratifies cortical areas along spatial gradients of cortical microstructure and connectivity (Margulies *et al*., 2016; Huntenburg *et al*., 2018, Paquola *et al*., 2019*b*). Cortical areas indeed show variable microstructural characteristics, often following sensory-fugal spatial gradients that relate to plasticity and neural excitability (Deco *et al*., 2013; Wang and Knösche, 2013; Burt *et al*., 2018, Paquola *et al*., 2019*a*, *b*; Wang, 2020; Weng *et al*., 2020). For example, paralimbic circuits differ from sensory networks by having an agranular architecture with only subtle laminar differentiation and relatively low myelin content, while sensorimotor areas have a marked layer IV and higher intracortical myelin (Flechsig Of Leipsic, 1901; Barbas and García-Cabezas, 2015; Palomero-Gallagher and Zilles, 2019; Drenthen *et al*., 2020; Weng *et al*., 2020). Complementing these microstructural variations, recent work has shown gradients of functional connectivity running from sensorimotor networks towards heteromodal systems, notably the default mode network (Margulies *et al*., 2016). Contextualizing atrophy and atypical asymmetry patterns along these microstructural and functional connectivity gradients may shed light on potential anatomical determinants of cortical pathology in TLE.

We used the ENIGMA-Epilepsy dataset to map patterns of atypical inter-hemispheric asymmetry and regional atrophy in 732 individuals with TLE and 1,418 healthy controls. We systematically assessed the commonalities and divergences of these patterns, and contextualized findings with respect to microstructural and functional connectivity gradients, derived from parallel myelin-sensitive microstructural MRI and resting-state functional acquisitions (Margulies *et al*., 2016, Paquola *et al*., 2019*b*; Vos de Wael *et al*., 2020). We formulated the following hypotheses: (i) the spatial distribution of TLE-related cortical asymmetry and atrophy would differ, with the former being particularly temporo-limbic; (ii) atypical asymmetry and atrophy maps would relate to cortical gradients, with the asymmetry map being more closely related to the primary temporo-limbic gradients derived from cortical microstructure. We also assessed whether inter-hemispheric asymmetry and regional atrophy mapping would show differential associations with clinical parameters, notably effects of disease duration and age of onset. In addition to benefiting from the high power of ENIGMA-Epilepsy, we validated the consistency of our findings at the level of single patients and individual sites.

## Methods

### Participants

We analyzed 2,150 T1-weighted MRI datasets from 732 patients with TLE and confirmed/suspected mesiotemporal sclerosis (55% females, mean age ± SD = 38.56 ± 10.61 years; 391/341 left/right TLE) and 1,418 healthy controls (55% females, mean age ± SD = 33.76 ± 10.54 years) obtained from 19 different sites via the Epilepsy Working Group of ENIGMA (Whelan *et al*., 2018, Larivière *et al*., 2020*c*; Sisodiya *et al*., 2020) (**Table 1**). Individuals with epilepsy were diagnosed by epilepsy specialists at each center according to classifications of the International League Against Epilepsy (ILAE) (Berg *et al*., 2010). TLE patients were diagnosed based on electroclinical and neuroimaging findings. Participants with a primary progressive disease (*e.g*., Rasmussen’s encephalitis), visible malformations of cortical development, or prior neurosurgery were excluded. For each site, local institutional review boards and ethics committees approved each included cohort study, and written informed consent was provided according to local requirements.

**Table 1.** Demographic information of individuals with TLE and site-matched controls. Means and SDs are reported.

| Information | ENIGMA-Epilepsy |  | HCP (HC) | MICs |  |
| --- | --- | --- | --- | --- | --- |
|  | TLE | HC |  | TLE | HC |
| Number | 732 | 1,418 | 207 | 23 | 53 |
| Age (years) | 38.56 ± 10.61 | 33.76 ± 10.54 | 28.73 ± 3.73 | 37.29 ± 11.96 | 30.84 ± 7.59 |
| Sex<br>(male:female) | 329:403 | 643:775 | 83:124 | 11:12 | 33:20 |
| Age at onset<br>(years) | 16.07 ± 12.27 | N/A | N/A | 21.59 ± 11.65 | N/A |
| Side of focus<br>(left/right) | 391/341 | N/A | N/A | 15/7<br>(1 bilateral) | N/A |
| Duration of<br>illness (years) | 22.74 ± 14.06* | N/A | N/A | 15.82 ± 12.45 | N/A |

\*Information available in 695 TLE patients.
*Abbreviations:* SD, standard deviation; TLE, temporal lobe epilepsy; HC, healthy control; HCP, Human Connectome Project; MICs, microstructure-informed connectomics; N/A, not available.

Gradients were derived from two independent cohorts containing healthy controls and patients with TLE: (i) A sample of 207 unrelated healthy young adults (60% females, mean age ± SD = 28.73 ± 3.73 years) from the HCP (Van Essen *et al*., 2013), (ii) A sample of 53 healthy controls (38% females, mean age ± SD = 30.84 ± 7.59 years) and 23 TLE patients (52% females, mean age ± SD = 37.29 ± 11.96 years) from our local site at the MNI (microstructure-informed connectomics; MICs). All participants gave written and informed consent.

### Data preprocessing

#### a) ENIGMA data

Participants underwent T1-weighted scans at each of the 19 centers, with acquisition protocols detailed elsewhere (Whelan *et al*., 2018). Imaging data were processed by each center through the standard ENIGMA workflow. In brief, individual surface modeling was performed using FreeSurfer 5.3.0 (Dale *et al*., 1999, Fischl *et al*., 1999*a*, *b*; Fischl, 2012), including magnetic field inhomogeneity correction, non-brain tissue removal, intensity normalization, and segmentation. White and pial surfaces were fit along tissue boundaries. Surfaces were inflated to spheres, followed by spherical registration to standard (fsaverage) space. Based on the Desikan-Killiany atlas (Desikan *et al*., 2006), cortical thickness was measured across 68 grey matter brain regions.

#### b) HCP data

T1- and T2-weighted, as well as rs-fMRI data, were obtained using a Siemens Skyra 3T at Washington University (Van Essen *et al*., 2013). The T1-weighted images were acquired using a magnetization-prepared rapid gradient echo (MPRAGE) sequence (repetition time (TR) = 2,400 ms; echo time (TE) = 2.14 ms; inversion time (TI) = 1,000 ms; flip angle = 8°; field of view (FOV) = 224 × 224 mm^2^; voxel size = 0.7 mm isotropic; and number of slices = 256). T2-weighted data were obtained with a T2-SPACE sequence (TR = 3,200 ms; TE = 565 ms; flip angle = variable; FOV = 224 × 224 mm^2^; voxel size = 0.7 mm isotropic; and number of slices = 256). The rs-fMRI data were collected using a gradient-echo echo-planar imaging sequence (TR = 720 ms; TE = 33.1 ms; flip angle = 52°; FOV = 208 × 180 mm^2^; voxel size = 2 mm isotropic; number of slices = 72; and number of volumes = 1,200 per time series). During the rs-fMRI scan, participants were instructed to keep their eyes open, looking at a fixation cross. Two sessions of rs-fMRI data were acquired; each contained data of left-to-right and right-to-left phase-encoded directions, providing up to four time series per participant.

HCP data underwent minimal preprocessing pipelines using FSL, FreeSurfer, and Workbench, briefly summarized as follows (Fischl, 2012; Jenkinson *et al*., 2012; Glasser *et al*., 2013):

##### b-i) T1- and T2-weighted data

Data were corrected for gradient nonlinearity and b0 distortions, and the T1- and T2-weighted data were co-registered using a rigid-body transformation. The bias field was adjusted using the inverse intensities from the T1- and T2-weighting. White and pial surfaces were generated using FreeSurfer (Dale *et al*., 1999, Fischl *et al*., 1999*a*, *b*; Fischl, 2012). A midthickness surface was generated by averaging white and pial surfaces, and used to generate the inflated surface that was registered to the Conte69 template (Van Essen *et al*., 2012) using MSMAll (Glasser *et al*., 2016) and downsampled to a 32k vertex mesh.

##### b-ii) Microstructure data

HCP provides a myelin-sensitive proxy based on the ratio of the T1- and T2-weighted contrast (Glasser and Van Essen, 2011; Glasser *et al*., 2014). Here, we first generated 14 equivolumetric cortical surfaces within the cortex and sampled T1w/T2w ratio values along these surfaces (Paquola *et al*., 2019*b*). A microstructural similarity matrix was constructed by calculating linear correlations of cortical depth-dependent T1w/T2w intensity profiles between different Desikan-Killiany parcels (Desikan *et al*., 2006), controlling for the average whole-cortex intensity profile (Paquola *et al*., 2019*b*). The matrix was thresholded at zero and log-transformed (Paquola *et al*., 2019*b*). A group-average matrix was constructed by averaging matrices across participants.

##### b-iii) rs-fMRI data

Data were corrected for distortions and head motion, and were registered to the T1-weighted data and subsequently to MNI152 space. Magnetic field bias correction, skull removal, and intensity normalization were performed. Noise components attributed to head movement, white matter, cardiac pulsation, arterial, and large vein related contributions were removed using FMRIB’s ICA-based X-noiseifier (ICA-FIX) (Salimi-Khorshidi *et al*., 2014). Time series were mapped to the standard grayordinate space, with a cortical ribbon-constrained volume-to-surface mapping algorithm. The total mean of the time series of each left-to-right/right-to-left phase-encoded data was subtracted to adjust the discontinuity between the two datasets, and these were concatenated to form a single time series data. A functional connectivity matrix was constructed via linear correlations of the fMRI time series of different Desikan-Killiany atlas parcels (Desikan *et al*., 2006). Fisher’s r-to-z transformations rendered connectivity values more normally distributed (Thompson and Fransson, 2016), and we averaged the connectivity matrices across participants to construct a group-average functional connectome, also available via the ENIGMA Toolbox (https://github.com/MICA-MNI/ENIGMA) (Larivière *et al*., 2020*b*).

#### c) MICs data

Data were acquired on a Siemens Prisma 3T scanner. Acquisition parameters were similar to the HCP dataset (T1-weighted: TR = 2,300 ms; TE = 3.14 ms; TI = 900 ms; flip angle = 9°; FOV = 256 × 180 mm^2^; voxel size = 0.8 mm isotropic; and number of slices = 320; quantitative T1 (qT1): same as T1-weighted except for TR = 5,000 ms and TE = 2.9 ms; TI = 940 ms; flip angle 1 = 4°; flip angle 2 = 5°; rs-fMRI: TR = 600 ms; TE = 30 ms; flip angle = 52°; FOV = 240 × 240 mm^2^; voxel size = 3 mm isotropic; number of slices = 48; and number of volumes = 700). MICs data were preprocessed using micapipe (https://github.com/MICA-MNI/micapipe), which integrates AFNI, FSL, FreeSurfer, ANTs, and Workbench (Cox, 1996; Avants *et al*., 2011; Fischl, 2012; Jenkinson *et al*., 2012; Glasser *et al*., 2013).

##### c-i) T1-weighted data

Data were de-obliqued, reoriented, intensity non-uniformity corrected, and skull stripped. Models of the inner and outer cortical surfaces were generated using FreeSurfer (Dale *et al*., 1999, Fischl *et al*., 1999*a*, *b*; Fischl, 2012), and segmentation errors were manually corrected.

##### c-ii) qT1 data

After registering qT1 data to FreeSurfer space using a boundary-based registration (Greve and Fischl, 2009), we generated 14 equivolumetric intracortical surfaces and sampled qT1 intensity as *in vivo* proxies of depth-dependent cortical microstructure (Paquola *et al*., 2019*b*). The microstructural profile similarity matrix was constructed using the same procedures as for the HCP data.

##### c-iii) rs-fMRI data

We discarded the first five volumes, removed the skull, and corrected for head motion. Magnetic field inhomogeneity was corrected using topup with reversed phase-encoded data (Andersson *et al*., 2003). After applying a high-pass filter at 0.01 Hz, noise components attributed to head movement, white matter, cardiac pulsation, arterial, and large vein related contributions were removed using ICA-FIX (Salimi-Khorshidi *et al*., 2014). Preprocessed time series were mapped to the standard grayordinate space, with a cortical ribbon-constrained volume-to-surface mapping algorithm. After regressing out time series spikes, a functional connectivity matrix was constructed by calculating linear correlations of time series between different Desikan-Killiany parcels (Desikan *et al*., 2006). We applied Fisher’s r-to-z transformation to the individual functional connectivity matrix and averaged across participants to construct a group-average functional connectome.

### Atypical inter-hemispheric cortical asymmetry and regional atrophy

We calculated inter-hemispheric asymmetry of cortical thickness using the following formula: *AI = (ipsi-contra) / |(ipsi+contra)/2|* (Bernasconi *et al*., 2003; Kong *et al*., 2018; Sarica *et al*., 2018), where *AI* is asymmetry index and *ipsi* and *contra* are the cortical thickness of ipsilateral and contralateral areas, respectively. The asymmetry index was z-normalized relative to site-matched pooled controls and sorted into ipsilateral/contralateral to the focus (Liu *et al*., 2016). It was then harmonized across different sites by adjusting for age, sex, and intracranial volume using ComBat, a batch-effect correction tool that uses a Bayesian framework to improve the stability of the parameter estimates (Johnson *et al*., 2007; Fortin *et al*., 2018). We compared the harmonized asymmetry index between individuals with TLE and controls using a general linear model implemented in SurfStat (Worsley *et al*., 2009). Multiple comparisons across brain regions were corrected using the FDR procedure (Benjamini and Hochberg, 1995). In addition to parcel-wise analysis, we stratified asymmetry measures according to seven intrinsic functional communities (visual, somatomotor, dorsal attention, ventral attention, limbic, frontoparietal, and default mode) (Yeo *et al*., 2011) and lobes (frontal, parietal, temporal, occipital, cingulate, and insular cortex). In addition to the atypical asymmetry index, we assessed cortical atrophy in TLE patients relative to controls. The cortical thickness measures were z-normalized, flipped hemispheres of right TLE, and harmonized as for the asymmetry index. We compared the harmonized cortical thickness between the groups and the findings were multiple comparison corrected using FDR (Benjamini and Hochberg, 1995) as well as stratified according to functional communities and lobes.

### Association to gradients of cortical microstructure and function

We assessed topographic underpinnings of TLE-related asymmetry and atrophy through spatial correlation analysis with microstructural and functional gradients, the principal eigenvectors explaining spatial shifts in microstructural similarity and functional connectivity (Margulies *et al*., 2016, Paquola *et al*., 2019*b*). Gradients were defined using two alternative datasets, either based on both the HCP (*i.e*., healthy controls) or based on the MICs (*i.e*., healthy controls and TLE patients), using BrainSpace (https://github.com/MICA-MNI/BrainSpace) (Vos de Wael *et al*., 2020). Specifically, we calculated a parcel-to-parcel affinity matrix for each feature using a normalized angle kernel considering the top 10% entries for each parcel. As in prior work (Margulies *et al*., 2016; Huntenburg *et al*., 2017; Hong *et al*., 2019, 2020, Paquola *et al*., 2019*a*, Larivière *et al*., 2020*d*; Müller *et al*., 2020; Valk *et al*., 2020; Vos de Wael *et al*., 2020; Park *et al*., 2021), we opted for diffusion map embedding (Coifman and Lafon, 2006), a non-linear technique that is robust to noise and computationally efficient (Tenenbaum *et al*., 2000; von Luxburg, 2007). It is controlled by two parameters α and t, where α controls the influence of the density of sampling points on the manifold (α = 0, maximal influence; α = 1, no influence) and t scales eigenvalues of the diffusion operator. The parameters were set as α = 0.5 and t = 0 to retain the global relations between data points in the embedded space, following prior applications (Margulies *et al*., 2016; Hong *et al*., 2019, Paquola *et al*., 2019*a*, *b*; Park *et al*., 2020; Vos de Wael *et al*., 2020). We examined associations of the estimated gradients with cortical asymmetry in a single hemisphere and atrophy patterns in both hemispheres via linear correlations, where significance was determined using 1,000 non-parametric spin tests that account for spatial autocorrelation (Alexander-Bloch *et al*., 2018) implemented in the ENIGMA Toolbox (Larivière *et al*., 2020*b*).

### Consistency mapping across sites and individuals

We assessed the robustness of our findings within a probabilistic framework at the single site and subject level. The consistency across sites was measured by calculating linear correlations between epilepsy-related asymmetry and atrophy findings and gradients for each site. For individual-level consistency, we counted how many participants are comprised within a specific threshold (*i.e*., z < - 2). The counts were divided by the number of participants to obtain a probability map. Thus, the consistency probability indicates that the top N% patients showed extreme asymmetry or cortical atrophy measures in a given region. The consistency probability was correlated with microstructural and functional gradients, with 1,000 non-parametric spin tests (Alexander-Bloch *et al*., 2018, Larivière *et al*., 2020*b*).

### Associations with clinical variables

We associated clinical variables of duration and onset of epilepsy with atypical asymmetry index and cortical atrophy using supervised machine learning. We utilized five-fold nested cross-validation (Tenenbaum *et al*., 2000; Varma and Simon, 2006; Cawley and Talbot, 2010; Parvandeh *et al*., 2020) with least absolute shrinkage and selection operator (LASSO) regression (Tibshirani, 1996). We split the dataset into training (4/5) and test (1/5) partitions, and each training partition was further split into inner training and testing folds using another five-fold cross-validation. Within the inner fold, LASSO finds a set of non-redundant features (*i.e*., atypical asymmetry index or cortical atrophy of brain regions) that could explain the dependent variable (*i.e*., disease duration or onset age). Using a linear regression, we predicted the clinical variables of inner fold test data using the features of the selected brain regions. The model with best accuracy (*i.e*., minimum mean absolute error (MAE)) across the inner folds was applied to the test partition of the outer fold, and the clinical variables of outer fold test data were predicted. We repeated this procedure 100 times with different training and test partitions to avoid subject selection bias. We assessed the prediction accuracy by calculating linear correlations between the actual and predicted clinical variables with their 95% confidence interval across 100 repetitions, as well as MAE. The significance of the correlation between actual and predicted values was assessed using 1,000 permutation tests by randomly shuffling participant indices. A null distribution was constructed, and it was deemed significant if the real correlation value did not belong to 95% of the distribution (two-tailed p < 0.05). We compared our model with the baseline model (*i.e*., predicted clinical variable = mean(training set clinical variable)), and improved prediction performance of our model was assessed using Meng’s z-test (Meng *et al*., 1992). To assess whether the frequency of the selected brain regions derived from LASSO regression across cross-validations and repetitions is related to microstructural and functional gradients, we calculated spatial correlations between cortex-wide probability distributions and each of the gradients. Significance was assessed using 1,000 spin tests (Alexander-Bloch *et al*., 2018, Larivière *et al*., 2020*b*). As a post-hoc analysis, we correlated the cortical features of the highly probable regions (selected probability > 0.5) and clinical variables, and the significance was calculated based on 1,000 permutation tests by randomly shuffling participant indices.

### Sensitivity analysis

#### a) Left and right TLE

To assess whether left and right TLE show consistent results, we repeated assessing atypical cortical asymmetry and atrophy, and correlating the effects with gradients for separate left and right TLE subgroups.

#### b) Different density of connectivity matrix

In our main analysis, we estimated microstructural and functional gradients using connectivity matrices with 10% density. We repeated generating the gradients from connectivity matrices with different densities (20, 30, 40, 50%) and correlated with atypical cortical asymmetry and atrophy patterns.

#### c) Gradients generated using local dataset

To assess whether the gradients estimated using an independent dataset reveal consistent results with those derived from the HCP dataset, we generated microstructural and functional gradients using our local dataset, MICs, that included healthy controls, as well as individuals with TLE. We then computed the associations between cortical distortions and gradients.

#### d) Volumetric analysis

We additionally assessed atypical inter-hemispheric asymmetry and regional atrophy patterns of six subcortical regions (amygdala, caudate, nucleus accumbens, pallidum, putamen, thalamus), as well as the hippocampus, defined using Desikan-Killiany atlas (Desikan *et al*., 2006). We estimated the volume of each region, calculated asymmetry index (Bernasconi *et al*., 2003; Kong *et al*., 2018; Sarica *et al*., 2018), z-normalized both asymmetry index and volume of TLE patients relative to controls, flipped hemispheres of right TLE, and harmonized across different sites by adjusting for age, sex, and intracranial volume using ComBat (Johnson *et al*., 2007; Fortin *et al*., 2018). We compared asymmetry and regional volume between individuals with TLE and controls using a general linear model (Worsley *et al*., 2009). Next, we assessed the robustness of atypical asymmetry and atrophy by calculating consistency probability. Lastly, we performed the prediction analysis by considering both cortical thickness and subcortical/hippocampal volume measures using LASSO regression (Tibshirani, 1996) with five-fold nested cross-validation (Tenenbaum *et al*., 2000; Varma and Simon, 2006; Cawley and Talbot, 2010; Parvandeh *et al*., 2020). The prediction procedure was repeated 100 times with different training and test dataset, and the performance was measured using linear correlations between the actual and predicted clinical variables with their 95% confidence interval, as well as MAE. We compared our model with baseline model, and assessed improvement of the prediction performance using Meng’s z-test (Meng *et al*., 1992).

## RESULTS

### Atypical inter-hemispheric asymmetry patterns differ from regional cortical atrophy in TLE

We found significant deviations in inter-hemispheric asymmetry in TLE relative to controls, especially in lateral and medial temporal cortex, as well as precuneus, with ipsilateral regions being atypically smaller than contralateral regions (p_FDR_ < 0.05; **Fig. 1A**). Stratifying effects according to intrinsic functional communities (Yeo *et al*., 2011), highest deviations in asymmetry were observed in the limbic network followed by default mode and somatomotor networks (**Fig. 1B**). Lobar analysis identified most marked degrees of atypical asymmetry in the temporal lobes. Asymmetry patterns of TLE were markedly different from regional differences in bilateral cortical thickness. Indeed, comparing cortical thickness between TLE and healthy controls showed widespread and bilateral cortical thickness reductions in TLE, with strongest effects in precentral, paracentral, and superior temporal regions (p_FDR_ < 0.05; **Fig. 1A**). Findings were distributed across somatomotor, dorsal attention, and visual networks (**Fig. 1B**). Similarly, lobar stratification pointed to multi-lobar effects, most marked in frontal, parietal, and occipital lobes in both hemispheres. Notably, spatial correlations between atypical asymmetry and atrophy patterns in a single hemisphere were very low and did not surpass null models with similar autocorrelation (r = 0.05, p = 0.27) (Alexander-Bloch *et al*., 2018).

**Fig. 1.**
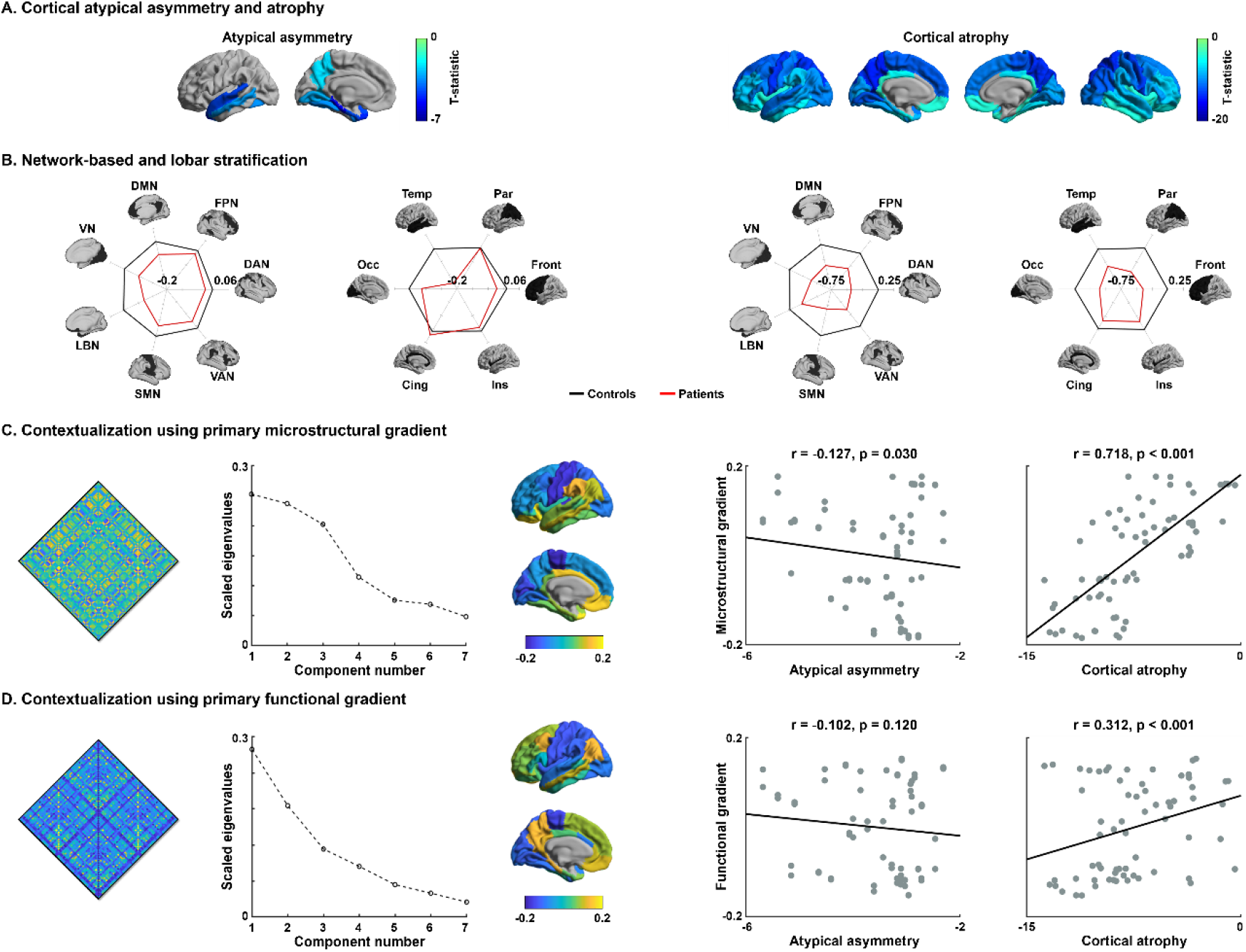
Topography of atypical cortical asymmetry and atrophy patterns in TLE. **(A)** Atypical inter-hemispheric asymmetry of cortical thickness and regional cortical atrophy between individuals with TLE relative to controls calculated using ENIGMA-Epilepsy dataset are shown on brain surfaces. (left) Blue regions indicate ipsilateral lateralization of cortical thickness or (right) decreases in cortical thickness in TLE relative to controls. **(B)** Effects (*i.e*., asymmetry index and cortical thickness) are stratified according to seven intrinsic functional communities (Yeo *et al*., 2011) and lobes. **(C)** Associations between epilepsy-related findings and microstructural/functional gradients calculated using HCP dataset. Cortex-wide microstructural profile similarity matrix and scree plot describing connectome variance after identification of principal eigenvectors are shown. The first principal eigenvector (microstructural gradient) is shown on the cortical surface. Spatial correlations between the principal microstructural gradient and TLE-related effects (*i.e*., atypical cortical asymmetry and atrophy) are reported with scatter plots. **(D)** Identical analysis to (C) but based on functional gradients. *Abbreviations*: VN, visual network; LBN, limbic network; SMN, somatomotor network; VAN, ventral attention network; DAN, dorsal attention network; FPN, frontoparietal control network; DMN, default mode network; Front, frontal; Par, parietal; Temp, temporal; Occ, occipital; Cing, cingulate; Ins, insular.

### A diverging topographic landscape of TLE-related atypical asymmetry and atrophy

Next, we assessed spatial associations of epilepsy-related findings with microstructural and functional gradients. The microstructural gradient depicted a continuous differentiation of cortical features between sensory and limbic areas, and was negatively correlated with atypical asymmetry index (r = −0.13, p_FDR_ = 0.03), reflecting elevated atypical asymmetry in temporo-limbic cortices in TLE (**Fig. 1C**). On the other hand, it was positively and markedly correlated with regional atrophy in TLE (r = 0.72; p_FDR_ < 0.001 **Fig. 1C**). The difference between these two correlations was significant (p < 0.001; Meng’s z-test) (Meng *et al*., 1992), indicating a dissociation of atypical asymmetry and atrophy patterns with respect to the primary microstructural gradient. The functional gradient differentiated primary sensory from transmodal regions, and did not show a significant association with atypical inter-hemispheric asymmetry in TLE (r = −0.10, p_FDR_ = 0.12), but a low-to-moderate positive association with regional atrophy (r = 0.31, p_FDR_ < 0.001) (**Fig. 1D)**.

### Consistency across sites and individuals

We confirmed the above topographic divergence across individual sites (**Fig. 2A**) by correlating microstructural and functional gradients with atypical asymmetry and regional atrophy in TLE for each site separately (**Fig. 2B**). These follow-up analyses confirmed our main findings (see *Fig. 2C*) that showed dissociation between atypical asymmetry and atrophy patterns. Multi-site analyses were expanded by assessing consistency at the level of individual patients (**Fig. 2C**). Prevalent atypical asymmetry was confirmed in somatomotor and limbic regions (**Fig. 2D**), and spatial patterns revealed significant associations only with the microstructural gradient (r = 0.13/-0.02, p_FDR_ = 0.04/0.42 for microstructural/functional gradients; **Fig. 2E**). The consistency probability of regional cortical atrophy showed higher consistency in sensory, precuneus, and temporal regions, and it showed significant negative correlations with both gradients (r = −0.29/-0.30, p_FDR_ <0.001/<0.001), supporting patient-level consistency.

**Fig. 2.**
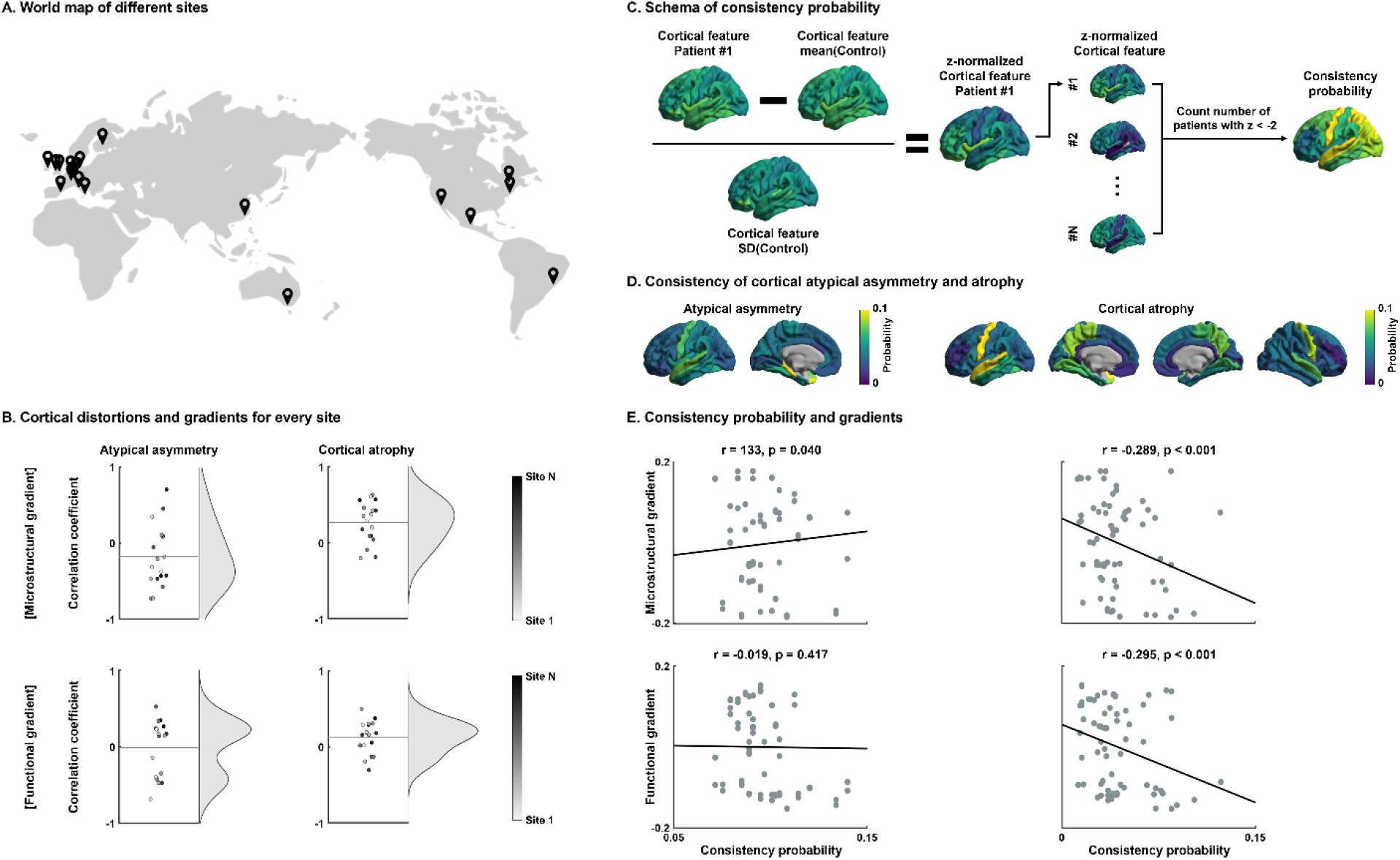
Consistency of atypical cortical asymmetry and atrophy. **(A)** World map of data acquisition sites. **(B)** Spatial correlations between topographic gradients and atypical cortical asymmetry/atrophy patterns of all sites. **(C)** Schema describing the computation of patient-wise consistency probability. The number of patients with large deviations of cortical features (*i.e*., atypical inter-hemispheric asymmetry or regional cortical atrophy) was counted. **(D)** Consistency probability of atypical cortical asymmetry and atrophy. **(E)** Spatial correlations between consistency probability and topographic gradients.

### Associations with clinical variables

Utilizing supervised machine learning, we probed associations of both atypical inter-hemispheric asymmetry and regional atrophy with disease duration and age at seizure onset. While cortical atrophy significantly predicted the clinical variables outperforming the baseline model (disease duration: mean ± SD r = 0.26 ± 0.02, MAE = 11.38 ± 0.10, Meng’s z-test p < 0.001; age at seizure onset: r = 0.17 ± 0.02, MAE = 9.91 ± 0.08, Meng’s z-test p = 0.01), atypical asymmetry did not (disease duration: Meng’s z-test p = 0.27; age at seizure onset: Meng’s z-test p = 0.20; **Fig. 3A, D**). Considering cortical atrophy, sensorimotor, medial/lateral temporal, and precuneus were frequently selected across cross-validations as the most important features used in the prediction for disease duration (**Fig. 3A**), and sensorimotor and limbic regions for age at seizure onset (**Fig. 3C**). As in the main analyses, we observed significant associations of the selected probability with connectome gradients (disease duration: r = −0.27/-0.34 p_FDR_ = 0.002/<0.001 for microstructural/functional gradients; age at seizure onset: r = −0.25/0.03 p_FDR_ <0.001/0.35; **Fig. 3B, E**). Associations in highly probable regions (selected probability > 0.5) were negative, *i.e*., disease duration/age at seizure onset associated with cortical thickness reductions (r = −0.30/-0.21, permutation-test p <0.001/<0.001; **Fig. 3C, F**).

**Fig. 3.**
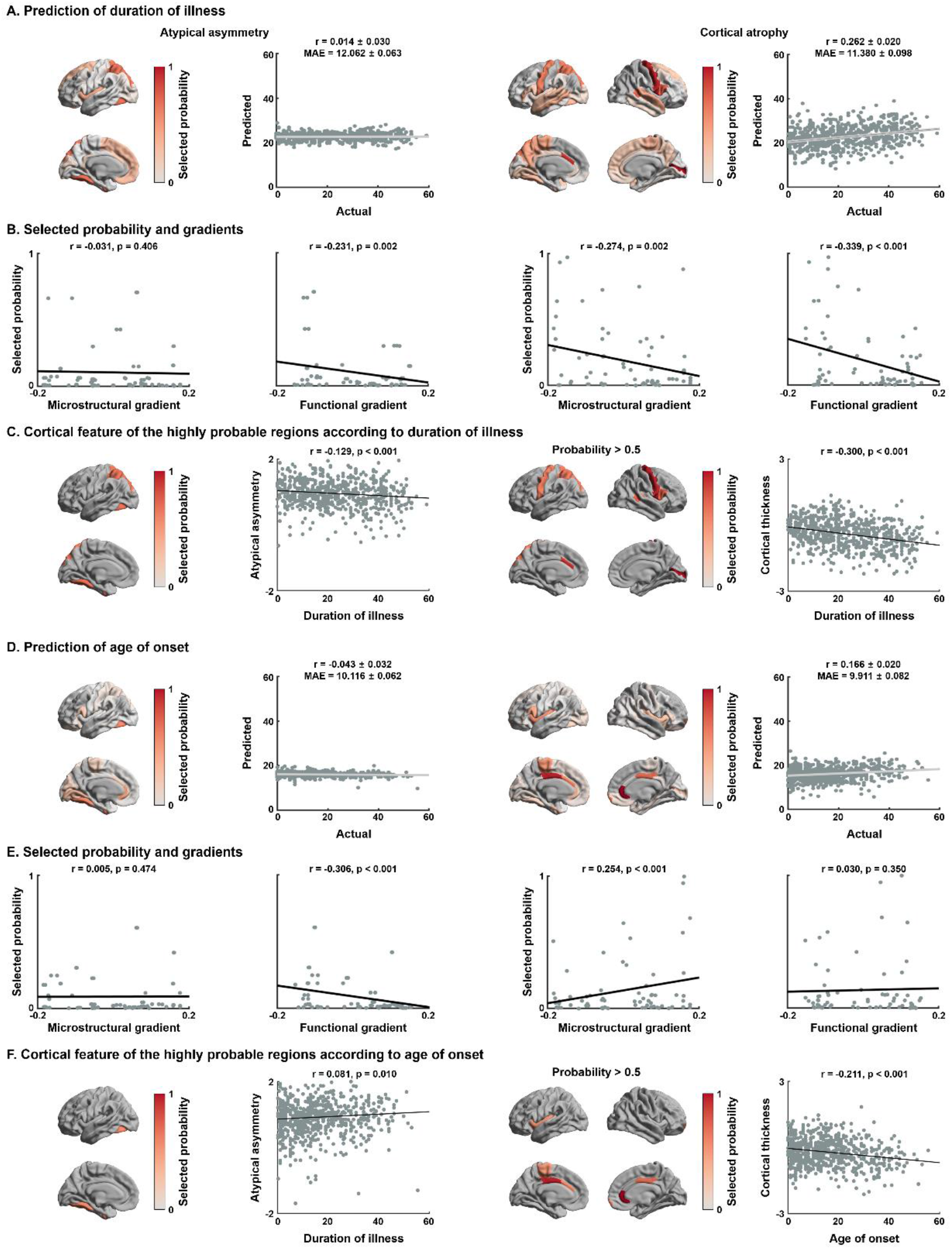
Associations between cortical features and clinical variables. **(A)** Probability of the selected brain regions across five-fold nested cross-validation and 100 repetitions for predicting duration of illness using atypical asymmetry index (left) and regional atrophy (right). Correlations between actual and predicted values of duration of illness are reported with scatter plots. Black lines indicate mean correlation, and gray lines represent the 95% confidence interval for 100 iterations with different training/test datasets. **(B)** Linear correlations between gradients and selected probability. **(C)** Spatial correlations between duration of illness and atypical asymmetry index (left), as well as cortical atrophy (right) in highly probable (selected probability > 0.5) regions. **(D-F)** Identical analysis to (A-C) but based on age at seizure onset. *Abbreviation*: MAE, mean absolute error.

### Sensitivity analyses

Several analyses supported robustness of our main findings.

#### a) Left and right TLE

We repeated the above analyses in left and right TLE separately. While the degree of asymmetry was stronger in left than right TLE, findings were overall consistent (**Fig. S1**).

#### b) Different density of connectivity matrix

We repeated our analyses by varying the thresholds of microstructural and functional connectivity matrices across different densities (20, 30, 40, 50%). Gradients and their associations with inter-hemispheric asymmetry, as well as regional atrophy, remained consistent (**Fig. S2**).

#### c) Gradients generated using local dataset

We also repeated the analysis after building gradients using a different dataset comprising both healthy individuals and patients with TLE. Microstructural and functional gradients were highly similar to those from the HCP dataset, and topographic associations were virtually identical and similarly robust as in the main findings (**Fig. S3**).

#### d) Volumetry of subcortical regions and the hippocampus

We also studied the volume of subcortical structures as well as the hippocampus. While atypical asymmetry and atrophy patterns both supported marked ipsilateral hippocampal effects (p_FDR_ < 0.05; **Fig. S4A**), spatial correlations between atypical inter-hemispheric asymmetry and regional atrophy were moderate and not significant (r = 0.51, p = 0.06). As for the cortical thickness-based results, these findings were consistent across individual subjects (**Fig. S4B**). When we considered both cortical thickness and subcortical/hippocampal volume, we were able to confirm our initial results in that atrophy but not atypical asymmetry related to age at seizure onset, while disease duration was significantly associated with both measures, outperforming the baseline model (atypical asymmetry: p = 0.004 for disease duration, p = 0.14 for age at seizure onset; atrophy: p < 0.001 for both disease duration and age at seizure onset; Meng’s z-test; **Fig. S4C**).

## Discussion

Together with the multi-site ENIGMA-Epilepsy initiative (Whelan *et al*., 2018, Larivière *et al*., 2020*c*; Sisodiya *et al*., 2020; Thompson *et al*., 2020), we investigated patterns of atypical inter-hemispheric asymmetry of cortical thickness and cross-sectional regional atrophy in a large sample of TLE patients and healthy controls. In particular, we studied whether (i) the spatial distribution of atypical inter-hemispheric asymmetry differed from patterns of regional atrophy in TLE relative to controls, (ii) these patterns follow different topographic principles of cortical organization, particularly with respect to microstructural and functional gradients, (iii) these effects showed a differential association to effects of epilepsy duration and age of onset. We found that atypical inter-hemispheric asymmetry analysis and regional atrophy mapping provide complementary insights into the pathology of TLE *in vivo*, with atypical asymmetry showing an ipsilateral limbic signature, while cross-sectional cortical thickness mapping indicated widespread and bilateral atrophy in TLE. Atypical asymmetry and atrophy patterns of the cortex were also differentially associated with microstructural and functional gradients representing core axes of cortical organization (Mesulam, 1998; Margulies *et al*., 2016; Burt *et al*., 2018; Huntenburg *et al*., 2018, Paquola *et al*., 2019*b*), supporting a topographic divergence of these two characterizations of TLE-related pathology. Findings were consistent across different sites and participants, corroborating generalizability. While cortical atrophy was correlated with disease duration and age at seizure onset, atypical asymmetry did not show an association to these variables. Collectively, our study underscores complementarity of atypical asymmetry and atrophy mapping for *in vivo* pathology mapping, which will be relevant for future imaging biomarker discovery and validation efforts.

In managing TLE patients, preoperative lateralization of temporal lobe pathology is key to define surgical target and often relies on the qualitative visual assessment of inter-hemispheric asymmetry. Quantitative imaging analyses in clinical and research settings can be geared towards the identification of asymmetry, and several prior studies have systematically investigated between-hemisphere differences in grey matter morphological measures in TLE. Most of these studies have focused on the asymmetry of the hippocampus and adjacent mesiotemporal structures, suggesting marked limbic structural asymmetry in TLE (Bernasconi *et al*., 2003; Bonilha *et al*., 2003, 2007; Shah *et al*., 2019). Asymmetry analysis has several benefits, including the ability to use a given patient as their own baseline while controlling for corresponding measures in controls. However, the field lacks systematic analyses of asymmetry, particularly outside the mesiotemporal region. There have been no quantitative comparisons of atypical inter-hemispheric asymmetry with maps of cross-sectional regional atrophy mapping, in which measures in patients with TLE are compared to groups of healthy controls. When carried out in structures of the limbic system, atrophy mapping also reveals structural compromise in TLE compared to healthy controls, with variable degrees of asymmetry ranging from relatively ipsilateral to rather bilateral depending on the TLE subgroups (Cendes *et al*., 1993; Bernasconi *et al*., 2003; Bernhardt *et al*., 2015). The advent of automated morphometric analysis has resulted in a predominance of studies focusing on cross-sectional regional thickness comparisons, and relatively few large-scale analyses have assessed the topography of atypical cortical thickness asymmetry patterns in TLE (McDonald *et al*., 2008, Bernhardt *et al*., 2009*a*; Alhusaini *et al*., 2012; Whelan *et al*., 2018; Galovic *et al*., 2019; Weng *et al*., 2020). Notably, although atypical asymmetry and atrophy are sometimes used interchangeably in the neuroimaging literature of TLE as *in vivo* indices of pathology, our findings pointed to differences in the patterns of atypical cortical asymmetry and regional patterns of cross-sectional atrophy in TLE. Spatial correlation analysis of their respective patterns confirmed this, failing to identify an association after accounting for spatial autocorrelations. Atypical asymmetry patterns in TLE followed a more specific paralimbic signature with maximal effects in the mesiotemporal lobe, in line with the classical conceptualizations of TLE as a limbic network disorder (Falconer *et al*., 1964; Margerison and Corsellis, 1966; Blanc *et al*., 2011; Tavakol *et al*., 2019). On the other hand, in line with prior single site analyses (Lin *et al*., 2007; Bernhardt *et al*., 2008, 2010; McDonald *et al*., 2008) and recent ENIGMA-Epilepsy studies (Whelan *et al*., 2018, Larivière *et al*., 2020*c*; Sisodiya *et al*., 2020), regional cortical atrophy mapping confirmed ipsilateral mesiotemporal atrophy in TLE, as well as widespread and bilateral effects outside paralimbic cortical areas. Findings were consistent in both left and right TLE patients. Thus, and despite both left and right TLE groups potentially showing different structural compromise (Bonilha *et al*., 2007; Coan *et al*., 2009; Santana *et al*., 2010; Kemmotsu *et al*., 2011; Dabbs *et al*., 2012; Liu *et al*., 2016; Whelan *et al*., 2018), findings overall suggest a similar divergence of atrophy and asymmetry patterns irrespective of seizure focus lateralization.

Our findings were further contextualized by quantifying the alignment of asymmetry and atrophy patterns along microstructural and functional gradients (Margulies *et al*., 2016; Huntenburg *et al*., 2018, Paquola *et al*., 2019*b*). Cortical microstructural gradients place sensorimotor cortices with strong laminar differentiation and high myelin content at one end and paralimbic regions with subtle myelination, low laminar differentiation, and increased synaptic densities at the other end (Huntenburg *et al*., 2017, Paquola *et al*., 2019*b*). Microstructural gradients, preserved across species (Huntenburg *et al*., 2017; Fulcher *et al*., 2019, Paquola *et al*., 2019*b*), follow canonical models of sensory-fugal cortical hierarchies (Mesulam, 1998), and capture inter-regional variations in heritability and plasticity (Vainik *et al*., 2020). While also starting at sensorimotor systems, the principal functional gradient radiates towards transmodal networks, such as the default mode and frontoparietal systems, and not the paralimbic cortices (Margulies *et al*., 2016). This divergence between microstructural and functional gradients may relate to less tethering of phylogenetically more recent association networks, such as the default mode network, from underlying signaling molecules (Buckner and Krienen, 2013) and may more closely reflect macroscale functional organization (Yeo *et al*., 2011). Spatial correlation analyses supported the dissociation of atypical cortical asymmetry and atrophy patterns with respect to microstructural gradients, where we observed increasing degrees of asymmetry towards the temporo-limbic anchor of the microstructural gradient, while atrophy patterns increased towards primary sensorimotor and unimodal association areas. While confirming stronger effects towards sensorimotor anchors in the case of atrophy patterns, functional gradient associations were less conclusive about atypical asymmetry, indirectly underscoring the paralimbic pattern of the latter. Furthermore, these findings may indicate that cortical morphological changes are better captured by microstructural than by functional hierarchies, a finding echoing prior associations between intracortical cellular-synaptic factors and measures of cortical thickness (Barbas and Pandya, 1989; Herculano-Houzel *et al*., 2013; Tomassy *et al*., 2014; Suminaite *et al*., 2019).

Big data initiatives such as ENIGMA-Epilepsy offer increased sensitivity to identify disease-related patterns of structural compromise and to assess consistency of findings at the single site and individual patient levels. We observed that the dissociation between atypical cortical asymmetry and atrophy remained consistent when we considered individual sites separately, and to some degree also at the level of individual participants. Using machine learning, we associated cortex-wide morphological data with clinical variables and showed inter-individual differences in cortical atrophy associated with disease duration and age at seizure onset. Associations were primarily driven by primary regions in sensorimotor cortex, together with temporal and precuneus regions. Unlike cortical thickness, atypical asymmetry patterns were not significantly associated with these clinical variables. These divergent clinical associations suggest that atypical inter-hemispheric asymmetry and regional cortical atrophy potentially reflect different TLE pathological processes, with asymmetry being more specifically related to an initial insult of the limbic circuitry. Alternatively, patterns of TLE-related atrophy in widespread and bilateral cortical territories had apparent progressive effects. The latter finding is consistent with prior cross-sectional, longitudinal, and meta-analytic findings assessing disease progression effects in TLE (Bonilha *et al*., 2006, Bernhardt *et al*., 2009*b*, 2010, 2013*b*; Coan *et al*., 2009; Caciagli *et al*., 2017; Galovic *et al*., 2019). This effect may relate to ongoing seizures, as supported by prior data showing associations to seizure frequency (Bonilha *et al*., 2006; Coan *et al*., 2009; Caciagli *et al*., 2017), as well as from anti-epileptic drug treatment (Pardoe *et al*., 2013; Caciagli *et al*., 2018). Moreover, drug-resistant patients are at increased risk for mood disorders and psychosocial challenges (Lin *et al*., 2012), which may furthermore adversely impact brain health.

We found that measures of atypical asymmetry and atrophy provide complementary windows into structural compromise in TLE, a finding also supported by the differential relationships to cortical topographic gradients and diverging associations to clinical parameters. Our findings advance our understanding of large-scale pathology in TLE and may direct future discovery and validation of clinically useful neuroimaging biomarkers.

## Acknowledgements

The authors would like to express their gratitude to the open science initiatives that made this work possible: (i) The ENIGMA-Epilepsy consortium and (ii) The Human Connectome Project (Principal Investigators: David Van Essen and Kamil Ugurbil; U54MH091657) funded by the 16 NIH Institutes and Centers that support the NIH Blueprint for Neuroscience Research; and by the McDonnell Center for Systems Neuroscience at Washington University. PS developed this work within the framework of the DINOGMI Department of Excellence of MIUR 2018-2022 (legge 232 del 2016).

## Supporting Information

**Fig. S1.**
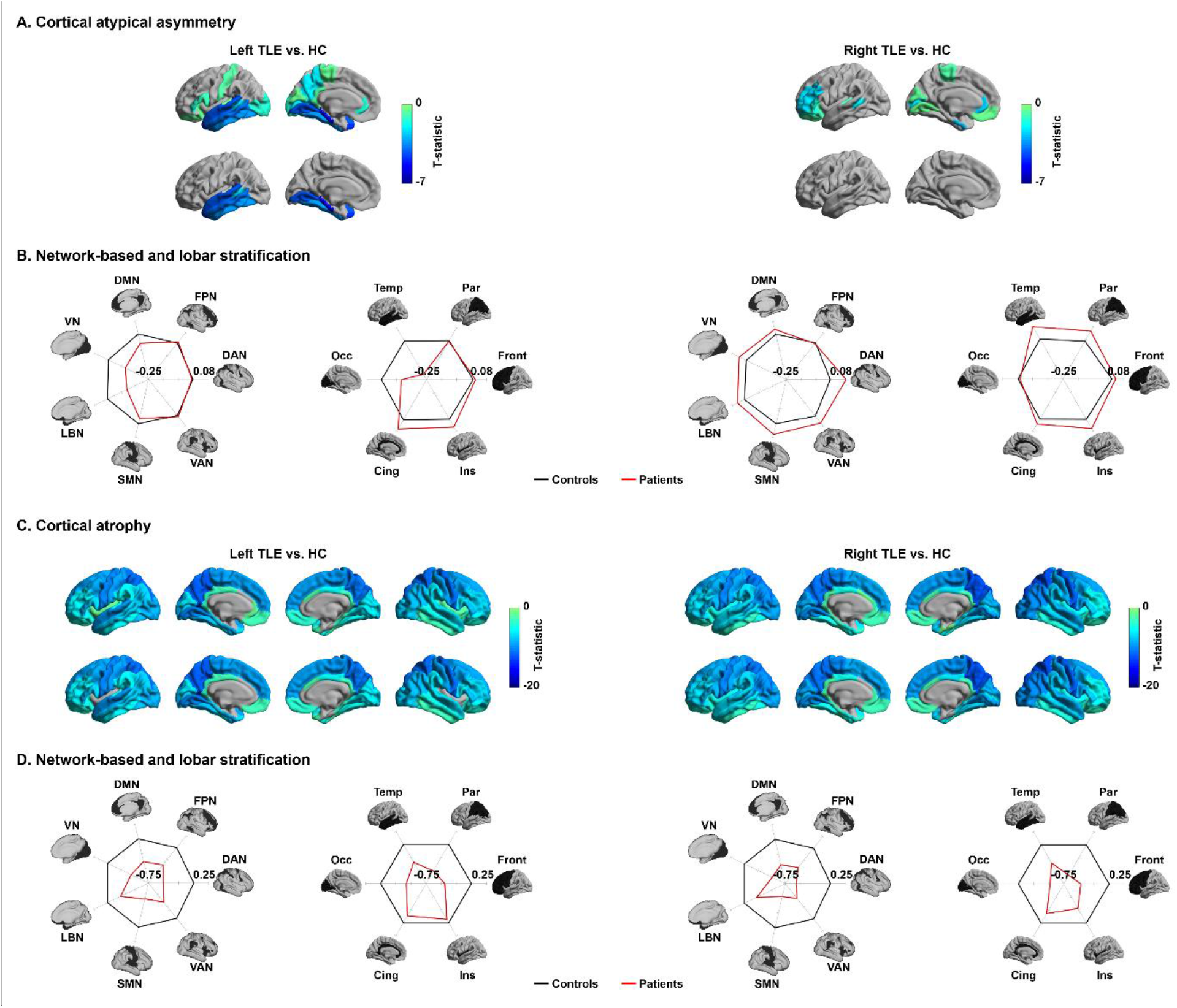
Results of left and right TLE. **(A)** Atypical cortical asymmetry in left and right TLE and **(B)** stratification of effects according to functional communities and lobes. **(C)** Associations of the effects with topographic gradients. **(D-F)** Results for cortical atrophy. For details, see *Fig. 1*.

**Fig. S2.**
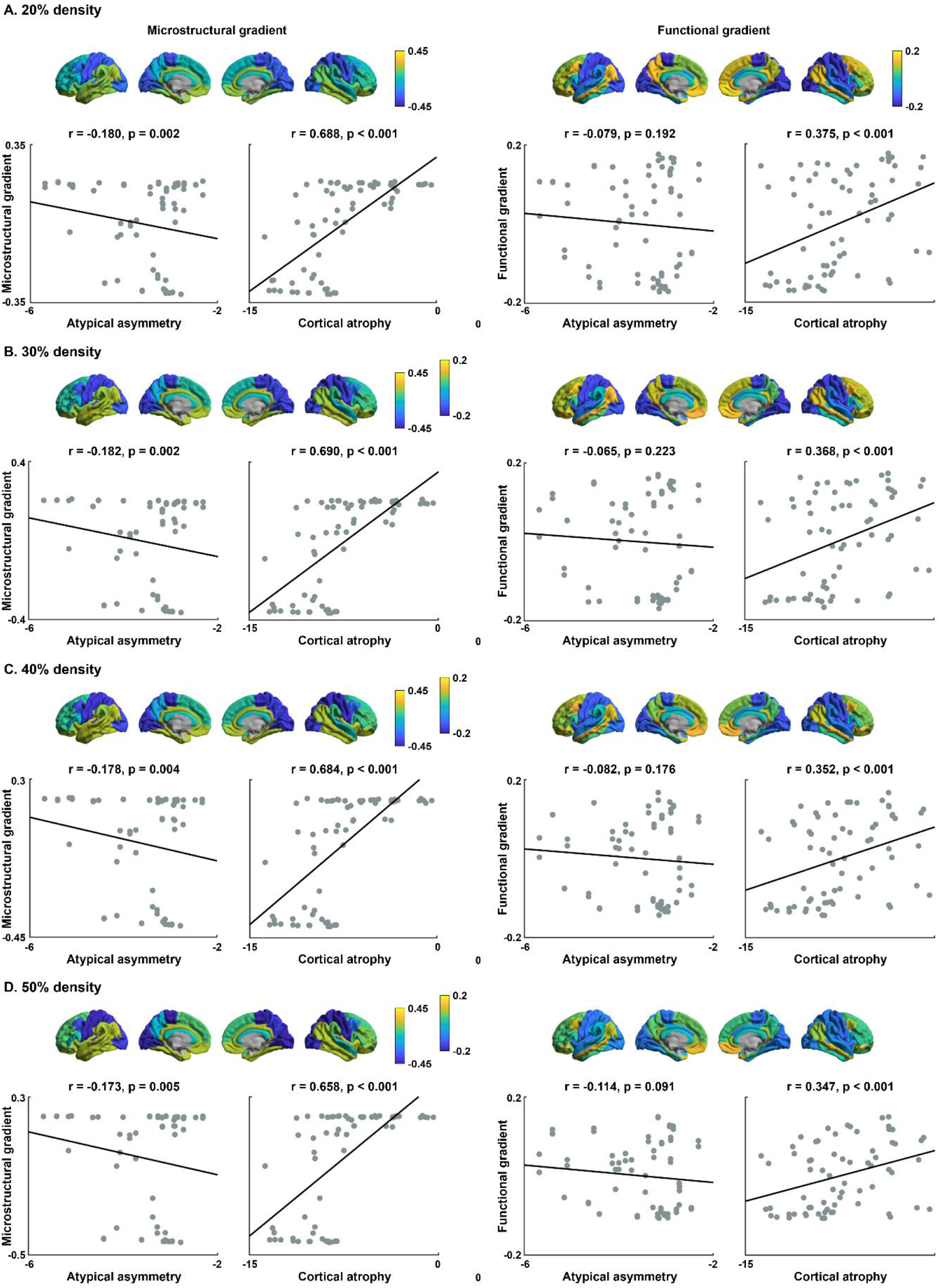
Results from different connectome densities. Microstructural and functional gradients derived from **(A)** 20, **(B)** 30, **(C)** 40, and **(D)** 50% density of connectivity matrices. Scatter plots show spatial correlations between microstructural/functional gradients and atypical cortical asymmetry (left) and atrophy (right).

**Fig. S3.**
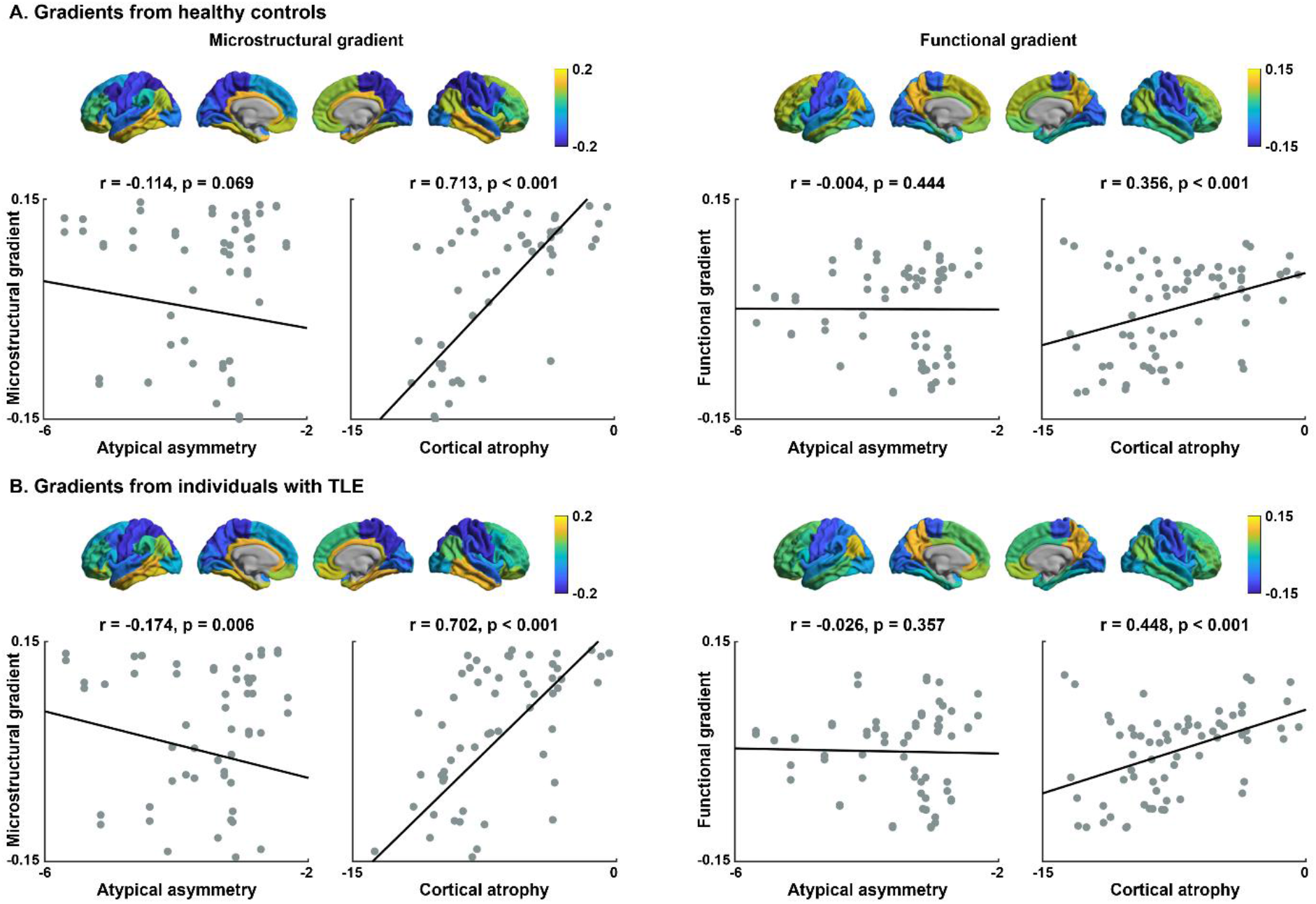
Gradients obtained from locally acquired microstructural and functional MRI data in healthy controls and patients with TLE. **(A)** Microstructural (left) and functional (right) gradients in healthy controls are shown on the brain surface. Associations between atypical cortical asymmetry and atrophy and these gradients are shown in the scatter plots. **(B)** Gradients and associations presented for patients with TLE.

**Fig. S4.**
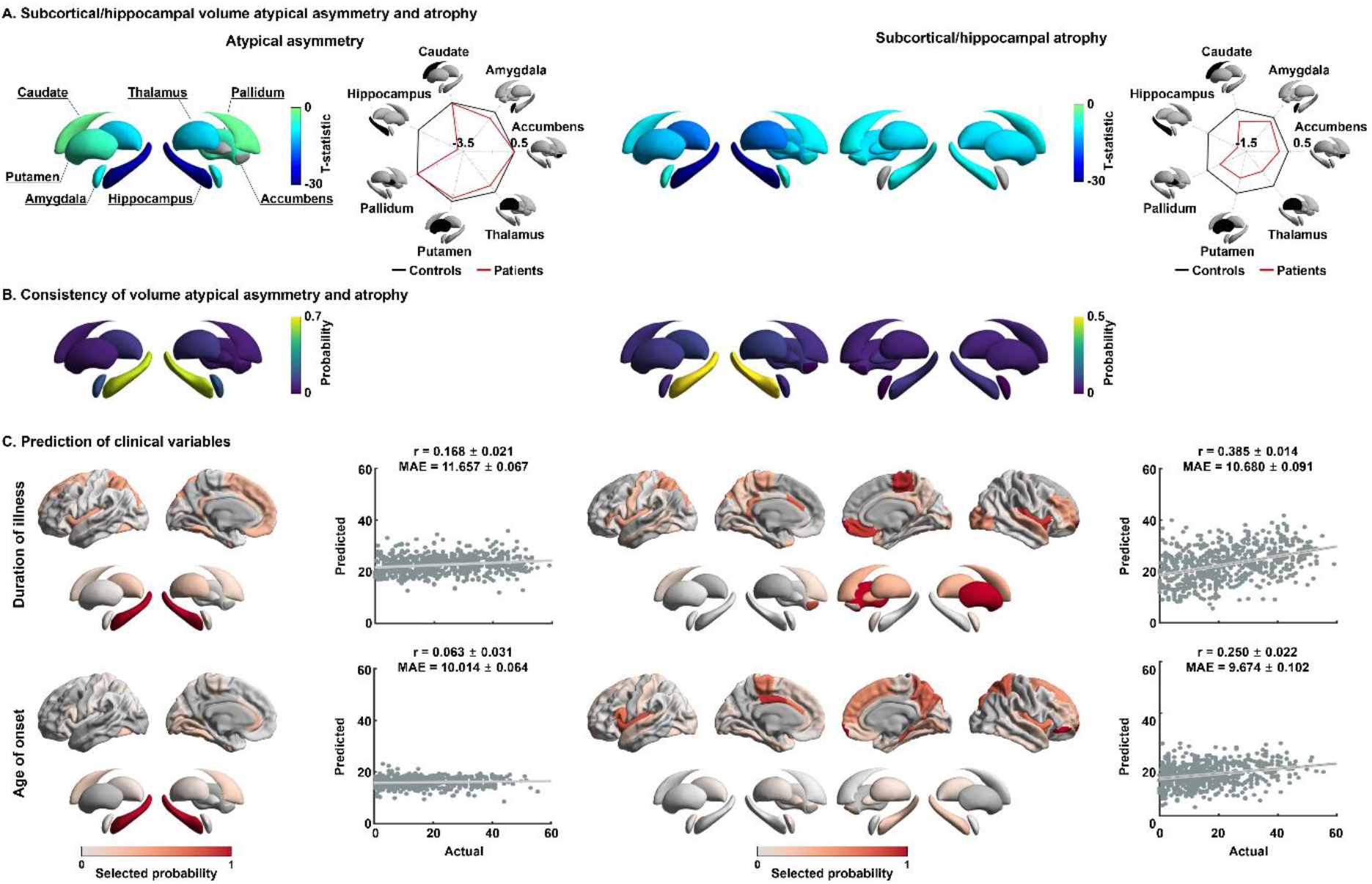
Analyses for subcortical and hippocampal volume. **(A)** Atypical asymmetry of subcortical/hippocampal volume differences and atrophy between individuals with TLE and controls. **(B)** Consistency probability of atypical asymmetry and atrophy in subcortical/hippocampal volume across individuals. **(C)** Prediction results for (top) duration of illness and (bottom) age at seizure onset using cortical and subcortical features. For details, see *Fig. 1, 2, and 3*.

